# Under-the-radar dengue virus infections in natural populations of *Aedes aegypti* mosquitoes

**DOI:** 10.1101/2020.01.24.919282

**Authors:** Sean M. Boyles, Carla N. Mavian, Esteban Finol, Maria Ukhanova, Caroline J. Stephenson, Gabriela Hamerlinck, Seokyoung Kang, Caleb Baumgartner, Mary Geesey, Israel Stinton, Kate Williams, Derrick K. Mathias, Mattia Prosperi, Volker Mai, Marco Salemi, Eva A. Buckner, John A. Lednicky, Adam R. Rivers, Rhoel R. Dinglasan

## Abstract

The incidence of locally acquired dengue infections increased during the last decade in the United States, compelling a sustained research effort on the dengue mosquito vector, *Aedes aegypti,* and its microbiome, which has been shown to influence virus transmission success. We examined the ‘metavirome’ of four populations of *Ae. aegypti* mosquitoes collected in 2016-2017 from Manatee County, Florida. Unexpectedly, we discovered that dengue virus serotype 4 (DENV4) was circulating in these mosquito populations, representing the first documented case of such a phenomenon in the absence of a local DENV4 human case in this county over a two-year period. We confirmed that all of the mosquito populations carried the same DENV4 strain, assembled its full genome, validated infection orthogonally by reverse transcriptase PCR, traced the virus origin, estimated the time period of its introduction to the Caribbean region, as well as explored the viral genetic signatures and mosquito-specific virome associations that potentially mediated DENV4 persistence in mosquitoes. We discuss the significance of prolonged maintenance of these DENV4 infections in *Ae. aegypti* that occurred in the absence of a DENV4 human index case in Manatee County with respect to the inability of current surveillance paradigms to detect mosquito vector infections prior to a potential local outbreak.

**Importance:** Since 1999, dengue outbreaks in the continental United States (U.S.) involving local transmission have occurred episodically and only in Florida and Texas. In Florida, these episodes appear to be coincident with increased introductions of dengue virus into the region through human travel and migration from endemic countries. To date, the U.S. public health response to dengue outbreaks is largely reactive, and implementation of comprehensive arbovirus surveillance in advance of predictable transmission seasons, which would enable proactive preventative efforts, remains unsupported. The significance of our finding is that it is the first documented report of non-outbreak DENV4 transmission and maintenance within a local mosquito vector population in the continental U.S.in the absence of a human case during a two-year time period. Our data suggest that molecular surveillance of mosquito populations in high-risk, high tourism areas of the U.S., may allow for proactive, targeted vector control before potential arbovirus outbreaks.

## Introduction

Approximately 40% of the globe is at risk of infection by flaviviruses, such as dengue virus (DENV): an enveloped, single-stranded RNA virus transmitted primarily by *Aedes aegypti* mosquitoes [1, 2]. Since severe disease from DENV infections can manifest as dengue hemorrhagic fever/dengue shock syndrome [1], DENV establishment in the continental United States is a major concern for public health agencies. In the USA, Florida has experienced increases in local DENV transmission since 2009 [3], driven in part by human and pathogen movement. *Ae. aegypti* is endemic throughout subtropical Florida and the vector population has resurged recently, following its near displacement by *Ae. albopictus* [4]. Autochthonous DENV infection occurs sporadically, primarily in Southern Florida with limited local cases elsewhere in the state [3]. In 2019, 16 cases of locally acquired DENV were reported for the state, including an area along the West-Central Florida Gulf Coast.

Recently, reports have indicated that certain insect-specific viruses (ISVs) can negatively impact or enhance arbovirus (including DENV) infections in insect cells [5, 6] and mosquitoes [7], respectively. Although the impacts of many ISVs on arboviral competence have yet to be determined, the evidence to date clearly indicates that the mosquito virome cannot be safely ignored and likely influences the risk of autochthonous DENV transmission once the virus is introduced into an area. Therefore, we conducted a metaviromic study of F_1_ (first-generation, lab-raised mosquitoes from wild parents) *Ae. aegypti* adult females collected as eggs from ovitraps in 2016-2017 from Manatee County to assess the potential risk of flavivirus transmission outside of Southern Florida. Although no indexed human case of DENV4 was reported during 2016-2017 in the county, we detected and sequenced DENV4, which may have been maintained vertically for at least one generation (but potentially more) in these *Ae. aegypti* mosquito populations along Florida’s Gulf Coast. We followed up this unexpected finding with genetic analyses to determine the DENV4 strain’s likely location of origin, assess the timeframe of virus introduction, and investigate strain-specific mutations that may have enabled adaptation to and/or persistence within local mosquito populations.

## Methods

### Mosquito sample preparation and viral RNASeq

*Ae. aegypti* eggs were collected in ovitraps in the summers of 2016-2017 (May 15, 2016, and June 19, 2017) from four Manatee County sites (**Fig. 1a**). To avoid cross contamination of mosquito viromes, each year eggs from each site were hatched independently in distilled water, reared to adulthood, speciated and then frozen. Female abdomens were pooled (N= 20/pool) separately for the four collection sites for a total of eight individual pools. Total RNA was extracted using the AllPrep DNA/RNA Mini Kit (Qiagen), and rRNA depleted with the NEBNext rRNA Depletion Kit (New England BioLabs). The NEBNext Ultra II Directional RNA Library Prep Kit (New England BioLabs) was used to prepare shotgun metagenomics libraries. Reverse-transcribed RNA libraries were sequenced using a HiSeq 3000 (Illumina) instrument in 2x101 run mode. The data were deposited into the NCBI Sequence Read Archive and Biosample archive under BioProject PRJNA547758.

**Figure 1.**
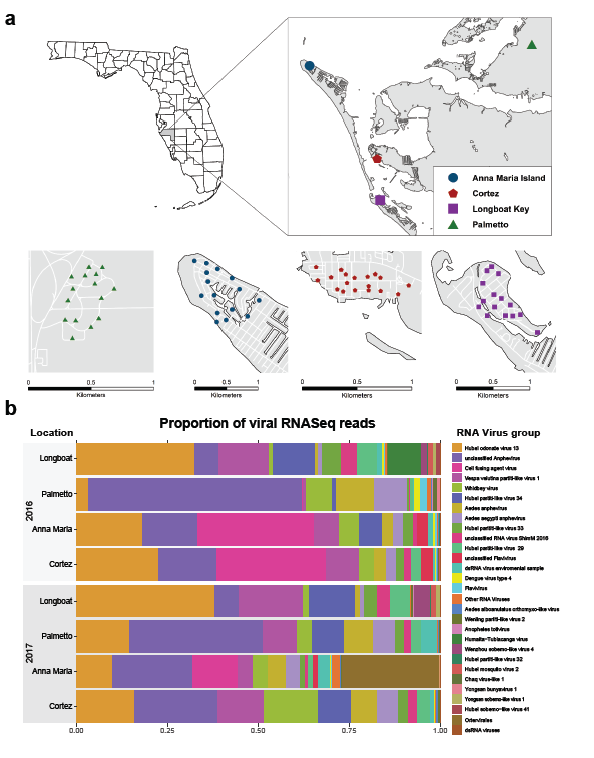
Metaviromic analysis of *Aedes aegypti* mosquito populations from Manatee County, Florida. **(a)** Locations of ovitraps in four different locations in Manatee County: Palmetto, Cortez, Ana Maria Island and Longboat Key. **(b)** The relative abundance of reads identified to come from RNA viruses in the 8 metagenomes. The proportion of the sub-composition is summarized at the species level for most viruses; however, some viruses were classified at higher levels if species could not be determined by the lowest common ancestor method.

### Initial assembly and metavirome analysis

BBduk (version 37.75; https://sourceforge.net/projects/bbmap/) was used to trim adaptor sequences and remove contaminants. *Ae. aegypti* sequences were removed using BBsplit (https://sourceforge.net/projects/bbmap/) against the *Ae. aegypti* Liverpool genome (AaegL5.1). Non-mosquito reads were assembled using Spades (3.11.1) in metagenomics mode [8]. For each contig, local similarity search in protein space was run using Diamond (0.9.17) [9] against the NCBI NR database. Reads were mapped against assemblies using Bowtie (2.3.4.1) [10], then sorted/indexed using Samtools (1.4.1) [11]. Megan 6 [12] was used to assign contigs and read counts to the lowest common ancestor (LCA) and to view viral contigs. To estimate microbial community abundance, Diamond (0.9.17) [9] was used to search reads against the NCBI NR database, Megan 6 [12] was used to assign read counts to the LCA, and R (3.6.0) package Compositions [13] (1.40-2) was used to create a sub-composition of RNA (**Fig. 1b**). Compositional count data from the Megan [12] LCA classification was assessed by ALDEx2 [14, 15] to estimate the statistical significance of the change in DENV4 reads from 2016 to 2017. ALDEx2 [14, 15] uses a Dirichlet multinomial Monte Carlo simulation to estimate the variance of the centered log ratio (CLR) values for taxa amongst the reads. Using the variance of the CLR, ALDEx2 [14, 15] computes P-values using Welch’s t-test and returns an effect size (CLR/variance) for the estimate. For a determination of the statistical significance of the observed decrease in CFAV reads from 2016-2017, a linear regression fitted to the CLRs of the Anna Maria and Cortez site DENV4 reads in 2016-2017 was utilized to yield an R^2^ value and a P-value to describe the trend.

### DENV4 refinement and genome-closing assembly

Two contigs covering most of the genome with a small gap were obtained. To create a closed genome, a dataset of genomes for DENV1-4 (NC_001477.1, NC_001474.2, NC_001475.2, NC_002640.1) and the two assembled contigs were used. We selected reads sharing a 31-mer with the dataset using BBduk (https://sourceforge.net/projects/bbmap/), followed by assembly with Spades in *meta* mode [8] and classification using Diamond [9] for a complete DENV4 genome. Read-mapping with Bowtie [10] revealed incorrect bases near the 3’ end, which were manually corrected. The genome was annotated using the Genome Annotation Transfer Utility [16] from the Virus Pathogen Database and Analysis Resource (ViPR) [17].

### Phylogenetic and Molecular Clock analyses

Two hundred thirty-four DENV4 genome sequences from GenBank (**Table S1**) were aligned using MAFFT version 7.407 [18] with the L-INS-I method [19]. IQ-TREE software [20] was used to evaluate phylogenetic signal in the genomes by likelihood mapping [21] and to infer maximum likelihood (ML) phylogeny based on the best-fit model according to the Bayesian Information Criterion (BIC) [20, 22]. Statistical robustness for internal branching order was assessed by Ultrafast Bootstrap (BB) Approximation (2,000 replicates), and strong statistical support was defined as BB>90% [23].

To estimate when DENV4 entered Florida, we used 145 strains including all isolates from the Americas, related Asian and African isolates, and randomly reduced oversampled Brazilian isolates. The strains in this dataset were not recombinant, as assessed by scanning the alignments for possible recombination points using the RDP, GENECONV, MaxChi, CHIMAERA, and 3Seq algorithms implemented in the RDP4 software (available from http://web.cbio.uct.ac.za/~darren/rdp.html) [24]. Correlation between root-to-tip genetic divergence and date of sampling was conducted [25] to assess clock signal before Bayesian phylodynamic analysis. Time-scaled trees were reconstructed using the Bayesian phylodynamic inference framework in BEAST v.1.8.4 [26, 27]. Markov Chain Monte Carlo (MCMC) samplers were run until 200/250 million generations to ensure Markov chain mixing, assessed by calculating the Effective Sampling Size (ESS) of parameter estimates. The HKY substitution model [28] was used with empirical base frequencies and gamma distribution of site-specific rate heterogeneity. The fit of strict vs. relaxed uncorrelated molecular clock models, and constant size vs. Bayesian Skyline Plot [29] demographic models were tested. Marginal likelihood estimates (MLE) for Bayesian model testing were obtained using path sampling (PS) and stepping-stone sampling (SS) methods [30, 31]. The best model was of a strict clock and constant demographic size. The maximum clade credibility tree was inferred from the posterior distribution of trees using TreeAnnotator specifying a burn-in of 10% and median node heights, then edited graphically in FigTree v1.4.4 (http://tree.bio.ed.ac.uk/software/figtree/), alongside ggtree available in R [32].

### Single-nucleotide variation analyses

The viral RNA sequencing reads were mapped onto the complete genome of seven DENV4 strains. These strains represent all the known DENV4 lineages (accession numbers are provided in **Fig. 3c**). We also mapped the reads onto the assembled Manatee DENV4 full genome. The read mapping was performed in the Geneious platform (Geneious Prime® version 2019.2.1) using the “map to reference” function under standard settings (Mapper: Geneious; Sensitivity: Highest Sensitivity/Slow; Fine tuning: Iterate up to 5 times; no trim before mapping). The Single nucleotide variation quantification was performed in the same platform using the “find Variation/SNV” function under default settings.

### DENV4 Genetic Analyses

From the 234 DENV4 genome alignment, sequences corresponding to the NS2A gene were extracted to investigate selection pressure and mutations that potentially influenced adaptation to and/or persistence in mosquito populations. Comparative selection and mutation analyses revealed NS2A as a relatively strong region of potential selection for the Manatee County genome. HyPhy algorithms were used to estimate non-synonymous (dN) to synonymous (dS) codon substitution rate ratios (ω), with ω<1 indicating purifying/negative selection and ω>1 indicating diversifying/positive selection [33, 34]. Fast, unconstrained Bayesian approximation (FUBAR) [35] was used for inferring pervasive selection, and the mixed effects model of evolution (MEME) [36] to identify episodic selection. Sites were considered to have experienced diversifying/positive or purifying/negative selective pressure based on posterior probability (PP) > 0.90 for FUBAR, and likelihood ratio test ≤ 0.05 for MEME.

To elucidate influential mutations in the Manatee DENV4 genome that potentially enabled persistence in the local mosquito population, a dN/dS analysis of the Manatee DENV4 against the relatively close, but geographically distant 1981 Senegalese DENV4 (MF004387.1) was conducted using JCoDA [37] with default settings, a 10-bp sliding window and a jump value of 5. To further assess the selective pressure throughout coding sequences in the DENV4 lineage that established transmission in Manatee County, we implemented a Single-Likelihood Ancestor Counting (SLAC) method [38] on the DataMonkey 2.0 web application [39]. It combines maximum-likelihood (ML) and counting approaches to infer nonsynonymous (dN) and synonymous (dS) substitution rates on a site-by-site basis for the different DENV4 coding alignments and corresponding DENV4 phylogeny. The measurements were performed on different alignments that included all strains, only genotype II strains, only clade IIa or IIb strains or only strains that are closely related to the DENV4 Manatee strain (multiple DENV4 coding sequence alignments are available as a Mendeley dataset). NS2A and 2K peptide genes were individually aligned and inspected between closely related DENV4 strains (1994 Haitian (JF262782.1), 2014 Haitian #1 (KP140942.1), 2014 Haitian #2 (KT276273.1), 2015 Haitian (MK514144.1), and 1981 Senegalese (MF004387.1) genomes) and the Manatee DENV4 for mutations to identify signals of adaptation of Manatee DENV4 to Floridian *Ae. aegypti*.

## Results

### DENV4 and ISVs in Ae. aegypti mosquitoes from Manatee County, Florida

Our metaviromic analysis of female *Ae. aegypti* mosquitoes detected DENV4 alongside several ISVs across four sites in 2016 and only Anna Maria and Cortez sites in 2017 (**Fig. 1a**). A full DENV4 genome (MN192436) was constructed with an overall genome coverage of ∼11X across the reads (**Fig. 2**). We observed that the 2017 DENV4 signal was much lower than 2016 for Anna Maria and Cortez (**Fig. S1**) and although Palmetto had the highest proportion of 2016 reads, this signal was virtually absent in 2017. To confirm DENV4 infection, we amplified and confirmed by direct sequencing the NS2A DENV4 amplicon for 2016 Longboat and Palmetto mosquito samples. Cumulatively, the drop in DENV4 relative to the metavirome from 2016-2017 was statistically significant (effect size=-2.026; P=0.035).

**Figure 2.**
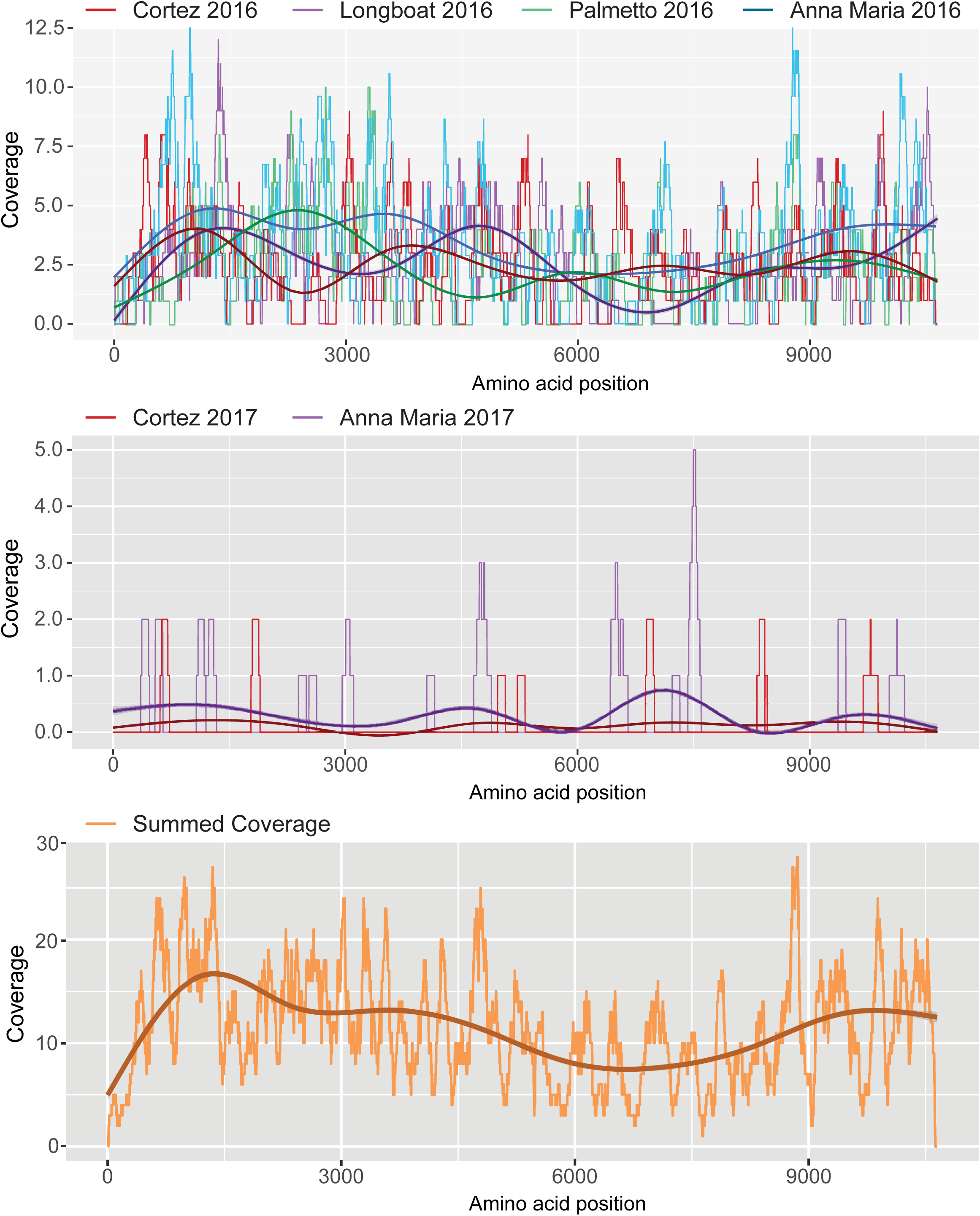
Mapping of RNASeq reads on the DENV4 genome. Coverage plots for DENV4 genome readings are shown from top to bottom graph panels. Coverage values across the genome for collection site/year combinations. Coverage is depicted on each y-axis and amino acid position on the x-axes. The smoothed central lines on the graphs indicate median values.

The RNA metavirome profile of the Manatee *Ae. aegypti* indicated an abundance of *Partitiviridae, Anphevirus,* Whidbey virus, and cell fusing agent virus (CFAV). *Partitiviridae* are known to primarily infect plants, protozoa, and fungi, but all the abundant groups in the metavirome have previously been detected in mosquitoes. We noted that the highest levels of CFAV (Anna Maria and Cortez sites) in 2016 were associated with DENV4 persistence into 2017 (P=0.07109; R^2^= 0.7943). Additionally, *Anphevirus* signals were notably abundant in the Palmetto samples in 2016 and 2017, coincident with DENV4 signal loss in Palmetto in 2017.

### DENV4 Phylogenetic and Molecular Clock Analyses

After analyzing the metavirome, we investigated the genome of the DENV4 strain to determine its likely source and assess the potential timeframe of introduction into Florida. Our first analysis confirmed the phylogenetic signal and absence of nucleotide substitution saturation (**Fig. S2a-b**). We subsequently explored Manatee County DENV4’s phylogeny with a 234-genome DENV4 dataset constructed from GenBank sequences (**Table S1**) by maximum likelihood (ML) phylogenetic inference (**Fig. 3a**). The ML phylogeny showed three clades: two Asian clades, and one American clade with two Senegalese strains (MF00438, KF907503) and one Thai (KM190936) at the base (**Fig. 3a**). Manatee DENV4 can be classified as DENV4 genotype IIb. The DENV4 genome obtained in Florida most closely clustered with two Haitian isolates from 2014 (KT276273, KP140942) and a cluster of Puerto Rican isolates (**Fig. 3a**). Further back, a Haitian isolate (JF262782) collected 20 years earlier also clustered with the Manatee-associated clade (**Fig. 3a**).

**Figure 3.**
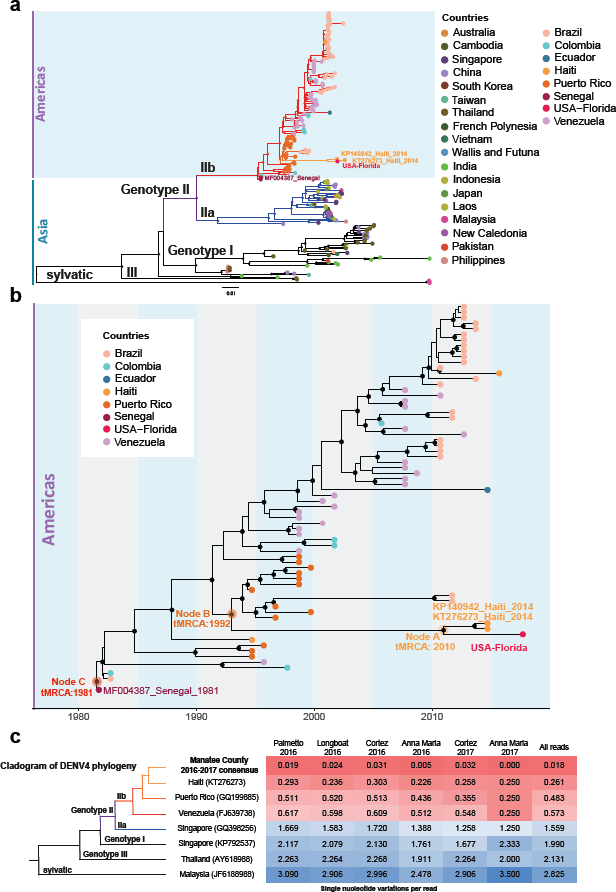
Phylogenetic and phylodynamic analyses of Manatee DENV4. **(a)** Maximum likelihood phylogenetic analysis of DENV4 full genome sequences. ML tree was obtained using IQ-TREE [20] software, diamonds indicate strong statistical support along the branches defined by ultrafast bootstrap >90. Tips are labeled and colored based on country of origin. **(b)** Bayesian phylodynamic reconstruction of DENV4 genotype IIb strains. The Maximum Clade Credibility time-scaled phylogenetic maximum clade credibility tree inferred using relaxed clock and constant demographic priors implemented in BEAST v1.8.4. Circles represent branches supported by posterior probability >0.90. Tips are colored based on location of origin. Labeled are nodes A (time of the most recent common ancestor [tMRCA] 2010), B (tMRCA 1992), and C (tMRCA 1981) on the branches. **(c)** SNVs/read per collection site/year combination of mosquitoes with significant detection by viral RNASeq in comparison to various reference genomes shown as a distance matrix is shown. The total numbers of SNVs were normalized by the total numbers of reads from each sample. Cell values refer to the SNV/read ratios of every sample (column) as compared to every representative sequence (rows). Cells are color-coded in the matrix as red = 0.0 SNV/read; white = 1.5 SNV/read; and blue = > 3 SNV/read.

To estimate the most recent common ancestor (MRCA) for DENV4 entry into Manatee County, Florida, as well as date divergence of the strain with Haitian isolates, we performed a molecular clock analysis using a Bayesian evolutionary framework [29] on a reduced dataset including only the “Americas clade.” We first assessed the phylogenetic signal and the absence of nucleotide substitution saturation (**Fig. S2c-d**) and then the temporal signal alone (**Fig. S3**). In the maximum clade credibility (MCC) tree, Manatee DENV4 clustered with the Haitian isolates from 2014 (Node A posterior probability [PP] > 0.9) (**Fig. 3b**). The MCC phylogeny showed that the time of the MRCA (tMRCA) for the DENV4 Manatee isolate and Haitian isolates was 2010 (Node A in **Fig. 3b**). This 95% high posterior density interval for this tMRCA suggests that DENV4 may have entered Manatee County sometime between 2006-2013. For Node B (**Fig. 3b**), the tMRCA of 1992 with a 95% HPD interval 1901-1994 indicated that Floridian and 2014 Haitian strains diverged from the 1994 Haitian DENV4 (JF262782), almost a decade before its arrival to Florida. However, strain divergence may have occurred in Haiti and was not necessarily precipitated by its introduction to Manatee County. Therefore, the introduction timeframe could be more recent than the estimated tMRCA.

### DENV4 SNVs/Read Analyses

Next, we examined the Manatee DENV4 genome sequences to compare strain variation between years and to identify mutations unique to the strain that potentially enabled local adaptation to and/or persistence in local mosquito populations. Following the MCC phylogenetic analysis, site-specific reads from mosquito populations in Manatee County were analyzed for single-nucleotide variations (SNVs) by the number of SNVs/read against the Manatee consensus genome and other global DENV4 genomes (**Fig. 3c**). SNV/read values showed only 22 SNVs across the 11,650-nucleotide Manatee County genome against all reads. SNVs were more substantial per read in the other DENV4 genomes. This indicates the likely persistence of a single strain of DENV4 in Manatee County during 2016-2017 transmission seasons.

### Signatures of Manatee DENV4 adaptation

We then explored selective pressures on the Manatee County DENV4 strain’s coding sequence that may be functionally important with respect to transmission and persistence of DENV4 in Floridian aegypti. The DENV4 genome has 5’ and 3’ untranslated regions (UTRs) flanking eight protein-coding genes (non-structural [NS] protein 1, NS2A, NS2B, NS3, NS4A, the 2K peptide, NS4B, and NS5) (**Fig. 4a**). The protein-coding regions of Manatee DENV4 were compared to four Haitian DENV4 genomes from 1994-2015 and a 1981 Senegalese DENV4 genome. These were analyzed for all amino acid substitutions between strains, and a dN/dS analysis was conducted comparing the Senegalese DENV4 genome with Manatee DENV4 (**Fig. 4a** and **Table S2**). The highest proportions of amino acid substitution were seen in NS2A and the 2K peptide; simultaneously, the highest dN/dS values occurred for the NS2A gene, to a point of weak positive selection (dN/dS > 1) that covered a V1238T mutation discussed further herein. We then calculated dN/dS ratios for DENV4 altogether, genotype II, genotype IIa, and genotype IIb with all sequences available, as well as within the Haiti-Florida clade and the Haiti-Florida-Puerto-Rico clades (**Fig. 4b**). Purifying selection, which occurs when non-synonymous mutations are deleterious, dominated, but we found weaker purifying selection in NS2A and 2K peptide genes, correlating to the Manatee-to-Senegal dN/dS analysis conducted previously. Values of dN/dS for these genes increased relative to those for flanking genes for genotype IIb and Caribbean/Florida-specific groups as well (**Fig. 4b**).

**Figure 4.**
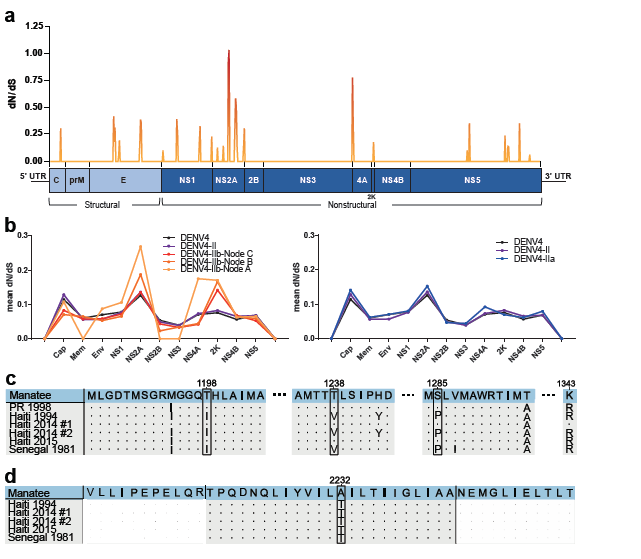
DENV4 amino acid analyses. **(a)** A dN/dS analysis conducted for the whole coding region of DENV4 Manatee vs. the Senegalese genome from 1981 (MF004387.1). The analysis was conducted utilizing full genome coding sequences in JCoDA using a sliding window analysis with a window size of 10. A small genome schematic is placed below the graph to its scale. Line colors approaching red from orange lie at higher values on the graph to indicate highe dN/dS values. **(b)** Using all available sequences, a dN/dS comparison was conducted to calculate mean ratio values within DENV4 overall, DENV4 genotype II, DENV4 genotype IIb, DENV4 FL-American-Caribbean clades, and DENV4-Florida-Caribbean-specific clades (respectively moving along Nodes C, B, and A from fig. 3B). Adjacent to the IIb-containing comparison, a comparison between all available DENV4 genome sequences, DENV4 genotype II, and DENV4 genotype IIa is depicted. **(c)** A comparative amino acid sequence alignment of Manatee County (MN192436), Puerto Rican (AH011951.2), Haitian 1994 (JF262782.1), Haitian 2014 #1 (KP140942.1), Haitian 2014 #2 (KT276273.1), Haitian 2015 (MK514144.1), and 1981 Senegalese (MF004387.1) genome sequences for the NS2A region sequenced in Puerto Rican isolate genomes by Bennet et al. [40]. Amino acid positions are numbered at the top of the figure. Key amino acid changes defining the 1998 DENV4 Puerto Rican outbreak in the NS2A gene are highlighted with boxes. **(d)** A comparison of the 2K peptide (colored in grey) sequence between the Manatee County (MN192436), 1994 Haitian (JF262782.1), Haitian 2014 #1 (KP140942.1), Haitian 2014 #2 (KT276273.1), Haitian 2015 (MK514144.1), and 1981 Senegalese (MF004387.1) genomes. Uncolored portions of the sequences correlate to portions of NS4A and NS4B.

Next, we further analyzed coding sequences in specific regions of the genome to investigate specific mutations that may have mediated Manatee DENV4 Floridian entry and persistence. The NS2A gene was analyzed in an alignment between Manatee (MN192436), 1994 Haitian (JF262782.1), 2014 Haitian #1 (KP140942.1), 2014 Haitian #2 (KT276273.1), 2015 Haitian (MK514144.1), and 1981 Senegalese (MF004387.1) genomes and a partial DENV4 genome (AH011951.2, Puerto Rico, 1998), with the analysis targeting three mutations that defined the 1998 DENV4 Puerto Rican outbreak [40] (**Fig. 4c**). The Manatee DENV4 sequence shares these key mutations with 1998 Puerto Rico, 2014 Haiti #1, and 2015 Haiti genomes. Conversely, the 1981 Senegal sequence and the oldest Haitian sequence from 1994 lack these mutations. In a selective pressure analysis utilizing the aforementioned 234-genome assembly, we observed strong background purifying selection with 143 sites that were found under episodic negative/purifying selection within the NS2A gene. Episodic diversifying/positive selection (evolutionarily preferred non-synonymous mutation) was detected in two sites corresponding to amino acids 1,238 and 1,333, both residues localized to transmembrane segments of the protein. This makes V1238T a mutation of note with the previous NS2A-associated analysis, detected in different analyses as a point of possible positive selection. The 2K peptide was next analyzed against the four Haitian genomes and the Senegalese genome from the first NS2A-specific analysis (**Fig. 4d**), and we observed that it had the second highest general rate of non-synonymous mutations and had a peak of weaker purifying selection (**Fig. 4a**). There was only one non-synonymous mutation among the six genomes, which is significant considering the size of the 2K peptide. This was a T2232A mutation present solely in the Manatee DENV4 sequence.

### DENV4 3’-UTR sequence and secondary structure analysis

To complete our genomic analysis of Manatee DENV4, we examined the 3’ UTR, as this region and its derivative subgenomic RNA have been implicated in epidemiologic and transmission fitness. Although the DENV4 3’ UTR lacks one of the two flaviviral nuclease-resistant RNAs (fNRs) in Domain I as compared to other DENV 3’ UTRs (3’ UTRs of other DENV serotypes [1–3]), DENV4 has the same conserved secondary structures in its domain II and III: two dumbbells (DB1 and DB2), and a 3′ end stem–loop (3′ SL) (**Fig. S4**). The 3’-UTR, through structural conformations, can affect viral replication in hosts [41]. We noted several transition substitutions in the DENV4 IIb lineage prior its arrival to Florida (Node B in **Fig. 3b** and **Fig. S4b**). Most of these mapped to either the highly variable region (HVR) or the adenine-rich segments that space functional RNA elements in DENV 3’ UTRs [42]. The U10318C substitution in fNR2 (fNR1 present only in other DENV serotypes) and the G10588A substitution on the 3’ SL mapped to base-pairing positions. However, these mutations have occurred in both directions in other lineages, suggesting they don’t imply fitness costs. Conversely, Floridian DENV4 underwent a rare transversion (A10478U) in a conserved position in DB2. This substitution favours formation of a new base-pair in DB2 structure. Additionally, an insertion (10467A) occurred in the adenine-rich segment upstream of DB2; this insertion is common for all lineages.

## Discussion

Our unbiased metavirome analysis of *Ae. aegypti* from Manatee County has revealed new insight into human arboviruses and ISV maintenance in a state prone to autochthonous flavivirus transmission. The observed drop in DENV4 relative to the mosquito virome (ISVs) between 2016-2017 was statistically significant (P=0.035), suggesting that the ISVs influence persistence of DENV4 in site-specific mosquito populations within the surveyed area. *Anphevirus* has been shown to reduce DENV viral titers *in vitro* during coinfections [43]. The abundance of Palmetto *Anphevirus* alongside the observed Palmetto 2016-2017 DENV4 reduction is consistent with this and suggests that these viruses and their respective abundance or relative proportions within a mosquito impact DENV4 prevalence in the vector population. The role of natural infections by insect-specific flaviruses on the proliferation of pathogenic arboviruses carried by different mosquito vector species is equivocal. A mosquito-specific flavivirus we detected known as cell fusing agent virus (CFAV) is of particular interest. Co-infection studies *in vitro* with DENV2 and CFAV result in enhanced proliferation in both [44]. Following this notion, the presence of CFAV in the same mosquito populations as DENV4 may improve viral dissemination and maintenance in mosquitoes. The observed correlation between persistence of DENV4 infection into 2017 in Anna Maria and Cortez mosquitoes with CFAV abundance in 2016 (**Fig. 1b**) appears to operate in parallel to the research conducted by Zhang et al. showing the enhanced replication of the two viruses [44]. An important caveat is that Zhang et al.’s research was conducted *in vitro*. Conversely, Baidaliuk, et al. demonstrated *in vivo* amplification-restrictive interaction between CFAV and DENV1 [7]. How DENV4 genotype X mosquito genotype X CFAV genotype interactions ultimately influence the vector competence of Floridian *Ae. aegypti* mosquitoes remains to be determined. The observed metavirome patterns sets the stage for follow-up studies to characterize the precise nature of ISV-DENV-mosquito interactions viz. vector competence.

The absence of an index human DENV4 case does not preclude the possibility that DENV4 was transmitted locally. Up to 88% of primary DENV infections are asymptomatic, with DENV4 being widely understood to cause primarily subclinical infections [45, 46]. Importantly, clinically inapparent infections could contribute to 84% of DENV transmission events through mosquitoes [45], so the threat of local transmission cannot be ruled out. However, it is noteworthy that DENV4 was detected in adult female mosquitoes reared from wild-captured eggs, implicating transovarial transmission (TOT) in local *Ae. aegypti* as has been shown for DENV1 in Key West, Florida [47]. However, since the DENV4 signal measured in 2017 was lower than in 2016, with two sites losing DENV4 prevalence, TOT alone may have been insufficient to maintain DENV4 from 2016-2017. Furthermore, we suspect that despite Manatee DENV4’s divergence from Haitian strains sometime between 2006-2013, it likely did not enter Manatee County until 2014 or after, given its similarity to the 2014-2015 Haitian DENV4 isolates and the fact that TOT is an inefficient process. Tertiary mechanisms, beyond ISV composition profile and TOT, could include inapparent human-mosquito infection cycles during the summer transmission (mosquito) season, which may have also contributed to DENV4 persistence in Manatee County aegypti. The exact mechanisms of maintenance in mosquitoes and proof of local transmission are difficult to elucidate at this juncture, considering all mosquito samples were processed for RNASeq and RT-PCR (i.e., no live virus can be isolated). Importantly, a comprehensive serosurvey with subsequent confirmation by gold-standard neutralization assay of the population from the four sample collection sites was not possible within the estimated mean half-life of detectable anti-DENV4 virion IgM or IgG. This limitation was unavoidable since (i) the complete viral genome assembly and orthogonal confirmation occurred more than two years following the initial mosquito collections, and (ii) there are significant confounders and logistical obstacles working with transient worker and migrant communities in the sampled area (well outside of the current scope of the study). However, the complete assembly and persistence over two years of an individual strain of DENV4, which is supported by results from orthogonal analytical approaches, remains provocative and reveals an unappreciated ecological process for DENV4 transmission in a non-endemic setting.

Tracking and predicting arbovirus movement and introduction into the United States, especially into Florida, can potentially lead to proactive efforts for increased monitoring and vector control at critical points of introduction into the state. DENV4 has been reported throughout the Caribbean, especially in Puerto Rico, Haiti and more recently in Cuba [48]. Florida has the largest populations of Puerto Rican, Haitian and Cuban origin and descent in the U.S., and there are ongoing efforts to develop effectively “sentinel” surveillance programs that can prepare Florida to deal with potential local arbovirus transmission. As expected, our analysis suggests a Caribbean origin for the Manatee isolate due to movements of DENV4 into Florida from Haiti, and preceding this, into Haiti from Puerto Rico. These results concur with previous findings depicting the Caribbean as a hotspot for arboviral spread in the Americas [48–50]. Diversifying selective pressure in the NS2A gene and the 2K peptide (**Fig. 4a-b**) experienced by American/Caribbean DENV4 may have contributed to the fixation of mutations driving the adaptation of DENV4 to environmental/vector conditions in these areas. NS2A mutations that characterized the 1998 DENV4 outbreak in Puerto Rico [40] are conserved between the Manatee, Puerto Rican, and two Haitian (JF262782.1 and KT276273.1) genomes (**Fig. 4c**). The 1981 Senegalese strain, the closest-clustering strain to the Manatee strain isolated outside the Americas (**Fig. 3a-b**), shares none of these mutations with Manatee DENV4. An in-depth understanding of how putative “hallmark” mutations in arboviruses can lead to increased local aegypti mosquito infections is lacking and compels further study.

We observed the expected 15-nucleotide deletion (Δ15) in the Manatee DENV4 3’ UTR (**Fig. S4**) that is present across all circulating DENV4 strains but absent from the extinct genotype I DENV4 lineage (GQ868594_Philippines_1956). Since the Δ15 deletion maps to the HVR, it does not alter the required secondary structures for sfRNA production. However, the HVR is an adenylate-rich unfolded spacer with poor sequence conservation—where no reliable secondary structure can be predicted, as our previous analyses suggested [42]. It has been speculated that these spacers favor the correct folding of adjacent functional structured RNA elements. The deletion might change the rate of folding of the downstream functional structured RNA and thus alter sfRNA production levels. Clearly, a closer molecular exploration of the exact role of this Δ15 deletion is needed.

The potential implications of our findings are profound; especially considering that arboviral surveillance of mosquito populations during the extended Florida mosquito season (April-October) is limited. To our knowledge, this is the first reported characterization of a DENV4 infection in native mosquito populations in Florida in the absence of an index human case across two years in a specific county. These data highlight the importance of knowing when and where arboviruses are introduced and point to the potential benefit of surveilling local mosquito populations for arbovirus infections prior to an outbreak. Given the increasing number of travel-related arbovirus introductions into Florida alone and the risk of local establishment in the state, we expect that while our report is seminal, it is likely the tip of the iceberg. If our data are any indication, the number of “under-the-radar” arbovirus infections of mosquito populations in migration hotspots across the state remains significantly underestimated.

## Acknowledgements

We gratefully acknowledge the support of Carina Blackmore, Danielle Stanek, and Andrea Morrison from the Florida Department of Health, as well as Lisa Conti, Kelly Friend, Davis Daiker, and Adriane Rogers from the Florida Department of Agriculture and Consumer Services for their institutional collaboration with the CDC Southeastern Center of Excellence in Vector Borne Diseases: The Gateway Program. We also thank Heather Coatsworth and Kaci McCoy for useful comments. This research was supported in part by the United States Centers for Disease Control (CDC) Grant 1U01CK000510-03: Southeastern Regional Center of Excellence in Vector-Borne Diseases: The Gateway Program. The CDC had no role in the design of the study, the collection, analysis, and interpretation of data, or in writing the manuscript. Support was also provided by the University of Florida Emerging Pathogens Institute, the University of Florida Preeminence Initiative, and United States Department of Agriculture, Agricultural research Service, project 6066-21310-005-00-D.

## Data Availability

Viral RNASeq read data is available in the NCBI Sequence Read Archive and Biosample archive under BioProject PRJNA547758. Genome sequence data for the Manatee sequence is available in NCBI’s GenBank database (MN192436) and reference sequences are available in the GenBank database with accession numbers described in the text. Multiple coding DENV4 sequence alignments from the dN/dS analyses and alignments for the RNA secondary structure model in Figure S4 are available with relevant accession numbers in a Mendeley dataset (https://data.mendeley.com/datasets/kwszjp63rb/draft?a=e11f9b80-bcfb-443b-918d-3016032ef3bd). The accession numbers in order from top to bottom for the compared sequences in Fig. 4c and 4d are MN192436, AH011951.2, JF262782.1, KP140942.1, KT276273.1, MK514144.1, and MF004387.1 (excluding AH011951.2 for Fig. 4d).

## Funding Statement

This research was supported in part by the CDC (https://www.cdc.gov/) Grant 1U01CK000510-03: Southeastern Regional Center of Excellence in Vector-Borne Diseases: The Gateway Program. The CDC had no role in the design of the study, the collection, analysis, and interpretation of data, or in writing the manuscript. Support was also provided by the University of Florida Emerging Pathogens Institute and the University of Florida Preeminence Initiative to RRD for this study. Mention of trade names or commercial products in this report is solely for the purpose of providing specific information and does not imply recommendation or endorsement by the U.S. Department of Agriculture.

Correspondence and requests for reprints should be sent to Dr. Rhoel R. Dinglasan. His e-mail address is. His FAX number is 352-392-9704 and his telephone number is 352-294-8448. His professional address is 2055 Mowry Road, 32611, Gainesville, FL.

---

**Figure S1. RNASeq read-proportion analyses.** To analyze the read abundance of the RNASeq assay conducted on the mosquito samples from Manatee county, the proportion of DENV4-mapped reads to the total number of reads mapped for each site and year pool was calculated. The number of reads that mapped specifically to DENV4 was divided by the total number of reads mapped, the resultant values being shown above the bars on the graph. Proportion values are on the y-axis of the graph.

**Figure S2 Assessment of phylogenetic quality for DENV4 strains. (a,c)** Phylogenetic signal, nucleotide substitution saturation and phylogenetic relationship in HIV envelope sequences from six patients obtained after ATI. Evaluation of the presence of phylogenetic signal satisfying resolved phylogenetic relationships among sequences was assessed by likelihood mapping (IQ-TREE: http://www.iqtree.org/), which estimates the likelihood of each of the three possible tree topologies for each group of four sequences (quartet) in the data set using the best-fit nucleotide substitution model chosen according to Bayesian Information Criterion (BIC). Quartets are considered “resolved” when the three likelihood are significantly different (phylogenetic signal), unresolved or partially resolved, when all three likelihood values or two of them are not significantly different (phylogenetic noise). Percentage within each triangle, indicate the proportion of resolved quartets (in the three corner areas), as well as the proportion of partially resolved (side areas) or unresolved (center) quartets. Extensive simulation studies have shown that side/center areas including <40% of the unresolved quartets can be considered robust in terms of phylogenetic signal [1, 2]. **(b,d)** Substitution saturation, which decreases the phylogenetic information contained in the sequences, was assessed using DAMBE7 (http://dambe.bio.uottawa.ca/DAMBE/) by plotting pairwise nucleotide (blue) transition (s) and (green) transversion (v) substitutions (y-axis) versus pairwise genetic distance (x-axis) determined with the Tamura and Nei 1993 (TN93) nucleotide substitution model [3].

**Figure S3. Assessment of temporal signal for DENV4 strains.** The plot represents regression analysis of root-to-tip genetic distance assessed using TempEst v1.5. The positive slope (R2=0.7135) indicates presence of temporal signal for the dataset.

**Figure S4. DENV4 3’ UTR Analyses. (a)** Alignment with the 3’-UTR of Manatee DENV4. The 3’-UTR RNA sequence alignment is of DENV4 from Manatee County, DENV4 Haiti 2014 #2 (KT276273.1), and DENV4 Philippines H241 (KR011349.2) genomes. The DENV4 RNA sequence alignment was generated with CLC Sequence Viewer 8.0 (https://www.qiagenbioinformatics.com/products/clc-main-w). Some DENV4 conserved 3’ UTR regions are designated in black boxes in the figure, including repeated conserved sequence 2 (RCS2), conserved sequence 1 (CS1), conserved sequence 2 (CS2), and 3’ upstream AUG region (3’ UAR). Different nucleotides are designated with different colors. **(b)** A diagram of the secondary structure of the DENV4 Manatee County 3’ UTR is depicted with key nucleotides and mutations highlighted and drawn in orange, correlating to nodes A or B from fig. 3B. Key secondary structure regions of the 3’ UTR are shown in black text: dumbbells 1-2 (DB1 & DB2), pseudoknots 1-5 (PK1-5), flavivirus nuclease-resistant RNA (fNR2), and the 3’ stem loop (3’ SL).

